# AutoGDC: A Python Package for DNA Methylation and Transcription Meta-Analyses

**DOI:** 10.1101/2024.04.14.589445

**Authors:** Chase Alan Brown, Jonathan D. Wren

**Affiliations:** University of Oklahoma Health Science Center, Oklahoma Medical Research Foundation; Oklahoma Medical Research Foundation

**Keywords:** Bioinformatics, Cancer, Genomic Data Commons, Database, Data analysis, Gene Expression, Epigenetics

## Abstract

**Motivation:** The Genomic Data Commons is a powerful resource which facilitates the exploration of molecular alterations across various diseases. However, utilizing this resource for meta-analysis requires many different tools to query, download, organize, and analyze the data. In order to facilitate a more rapid, simple means of analyzing DNA methylation and RNA sequencing datasets from the GDC we developed `autogdc`, a python package that integrates data curation and preprocessing with meta-analysis functionality into one simplified bioinformatic pipeline.

**Availability and Implementation:** The `autogdc` python package is available under the GPLv3 license at along with several examples of typical use-case scenarios in the form of a jupyter notebook. The data is all originally provided by the GDC, and is therefore available under the NIH Genomic Data Sharing (GDS) and NCI GDS policies.

## 1 Introduction

The Genomic Data Commons (GDC)^1^ is a rich and well-annotated human genomic data repository, containing many large studies on cancer with a focus on enabling discovery in precision medicine. The GDC has created several bioinformatic tools and web applications, which mostly operate via a representational state transfer (REST) application programming interface (API), in order to quickly query and analyze data. However, despite GDC’s work towards intuitive web apps and additional command line interface (CLI) tools, the python ecosystem for querying, downloading, processing, and analyzing GDC data in one python package is less developed. The `autogdc` package provides this access to the data repository, which contains more than 230,000 open-access data files and more than 350,000 controlled-access data files. The `autogdc` infrastructure focuses upon transcription and DNA methylation profiling data, as these two assay data types make up around 100,000 of the open-access files. The open-access paired DNA methylation and transcriptional profiling data provide unique opportunities for understanding molecular changes in cancer as well as in a broader biological context.

## 2 Similar work

### 2.1 GDC API and gdc - client

The Genomic Data Commons has produced several useful tools for increasing the accessibility of their data to bioinformaticians. One particular cli tool of interest is the `gdc-client` executable, which downloads data directly from the GDC servers given a manifest file. Due to the direct GDC support for this tool, which helps to standardize data transfer protocols from their servers, the `autogdc` package utilizes the gdc-client for constructing the large matrices that serve as the local backend for the different analyses. Additionally, the GDC has an API which is useful for gathering clinical annotations of the data, which is also wrapped within the `autogdc` package, therefore reducing much of the tedious data wrangling to match annotations and data.

### 2.2 TCGAbiolinks

The most functionally similar package to `autogdc` is TCGAbiolinks.^2^ However, TCGAbiolinks is implemented in R and does not include the sequence modeling tools for meta-analysis on the regulatory effect of DNA methylation on transcription. In contrast, `autogdc` is written for python developers and contains tools for answering various bioinformatic questions, including sequence modeling and elastic net aging models.

## 3 Implementation

The general architecture of the `autogdc` package is a ‘Dataset’ object, containing several methods to query, retrieve, and transform data from the GDC repository. This main object contains multiple data frames of genomic data, along with the corresponding annotation metadata for each sample. Additionally, in order to facilitate multi-omic studies, a ‘frame’ property can be called, which constructs a multi-indexed data frame of transcript and DNA methylation data, with information about DNA methylation loci, such as position to transcription start site, beta value, and it’s associated transcript.

This multi-indexed dataframe facilitates the rapid construction of summary statistics and tensors for building machine learning models to assess the epigenetic regulatory relationship with transcription.

### 3.1 DNA methylation and RNAseq data

All data is gathered automatically via `autogdc` from the Genomic Data Commons using the GDC API or the `gdc-client` command line interface executable. The compressed text files are then combined into dataframes of either DNA methylation or RNA sequencing count values, alongside the corresponding metadata from GDC. Additionally, preprocessing steps can be given as parameters, such that imputation of missing values and normalization will be automatically performed on the resulting matrices. Studies on regulation of transcription by DNA methylation were restricted to paired samples (within patient and tissue) with both 450k chip DNA methylation and RNA sequencing data. These restrictions yield a DNA methylation data frame consisting of 9472 samples and 396065 features and an RNA sequencing data frame consisting of 9415 samples and 59016 features. These matrices were processed by `autogdc` with mean imputation and quantile normalization for preprocessing.

### 3.2 Feature metadata

Infinium HumanMethylation450K loci meta data was retrieved from Illumina’s product file website and used to filter CpG sites via various genomic features. Additionally, Biomart^3^ was used to annotate gene symbols for RNA sequencing data.

### 3.3 Machine Learning Models

A long-short term memory recurrent neural network^4^ (tensorflow LSTM default implementation) with a latent state dimension of 32 units was used to encode the sequence of DNA methylation in order to predict RNA expression, as seen in Figure Figure 8. Several other models can be easily constructed in the package, such as transformer models to encode both RNA and DNA methylation states and predict tissues, age, etc.

## 4 Example Case Studies and Discussion

Several different `autogdc` case studies are presented in the jupyter notebook supplementary information, with a broader focus on the relation between transcription and DNA methylation.

### 4.1 Differential Gene Expression

One of the most common tasks for a meta-analysis is the determination of a set of differentially expressed genes between two contrasted groups. The `autogdc` package provides a simple interface to performing differential gene expression (See example cluster map of significantly altered genes in Figure 3) via the function `study.ddx(contrast = “sample_type”, method = “chdir”)`, which provides options for the method of analysis (characteristic direction^5^ or DESEq2^6^) or contrast variable. Simple gene set enrichments on top of the differential gene analysis are also provided via gseapy^7,8^.

**Figure 1.**
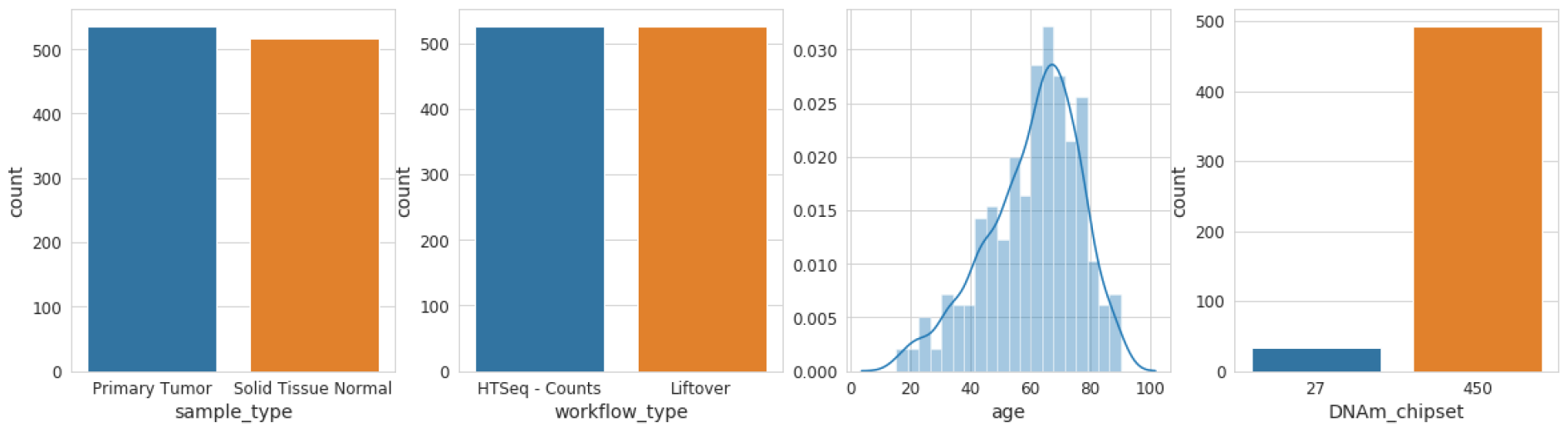
Summary Statistics of `autogdc` data

**Figure 2.**
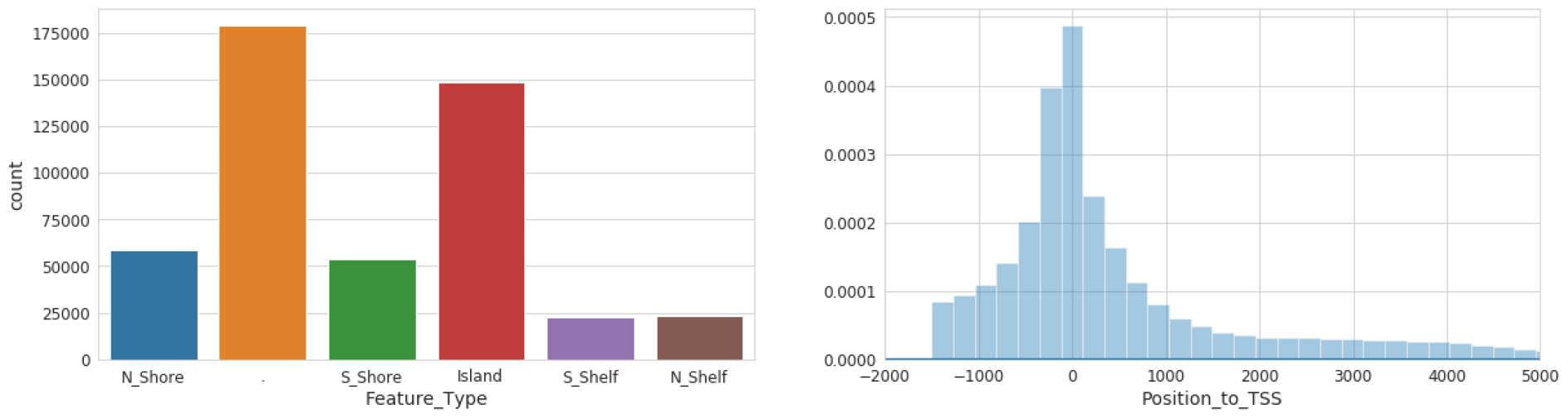
DNA Methylation loci distribution

**Figure 3.**
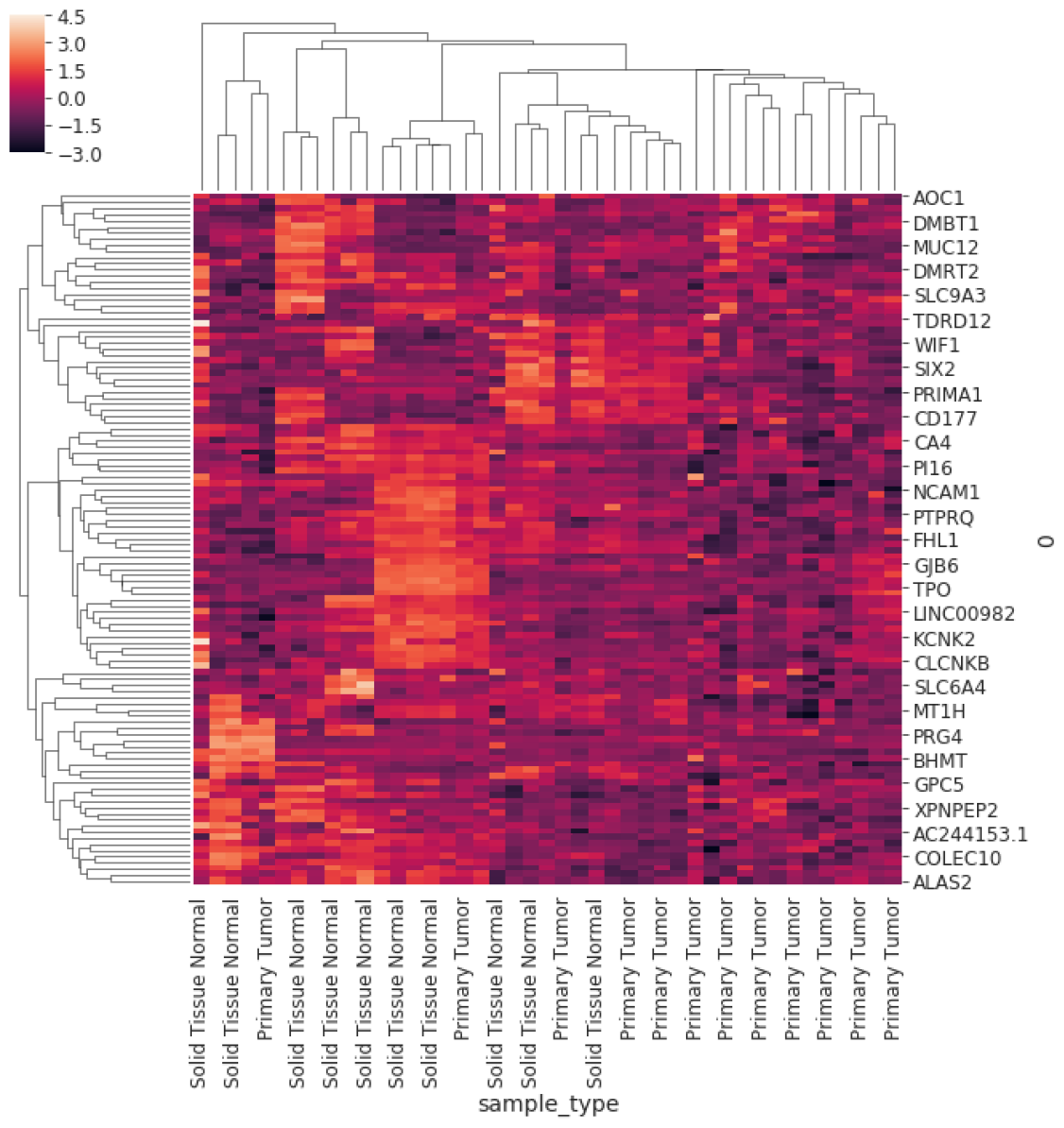
DNA Methylation loci distribution

### 4.2 DNA methylation-RNA expression relationship

Many studies have shown an inverse correlation between the average gene promoter DNA methylation and that gene’s level of mRNA expression^9^; however, the study of genomewide patterns has been less explored, mostly likely because of the lack of paired DNA and RNA datasets easily available to the public. AutoGDC enables this type of analysis to be done easily and quickly, as can be seen in the joint density plot of median promoter DNA methylation (loci within −1500 base pairs prior to the transcription start site) and gene expression for all genes (Figure 4). Interestingly however, when assessing the Pearson r correlation value between median DNA promoter methylation and RNA expression (natural logarithm of RNA sequencing counts) within each gene, the distribution of Pearson r correlation values contains a considerable number of positive correlations (Figure 5). In other words, the paired data from the GDC can be used to determine genes which have ‘noncanonical’ correlations between DNA methylation and transcription (high methylation and high expression or low methylation and low expression). A quick analysis of DNA methylation loci within promoters and the corresponding gene expression level shows several genes which have ‘non-canonical’ correlations (Figure 5). In addition to including functions to facilitate discovery with standard statistical techniques, `autogdc` contains several machine learning methods for studying the relationship between transcription and DNA methylation (Figure 8). These methods can be used to determine more detailed descriptions of genome-wide regulatory mechanisms and their dependence on sequence information.

**Figure 4.**
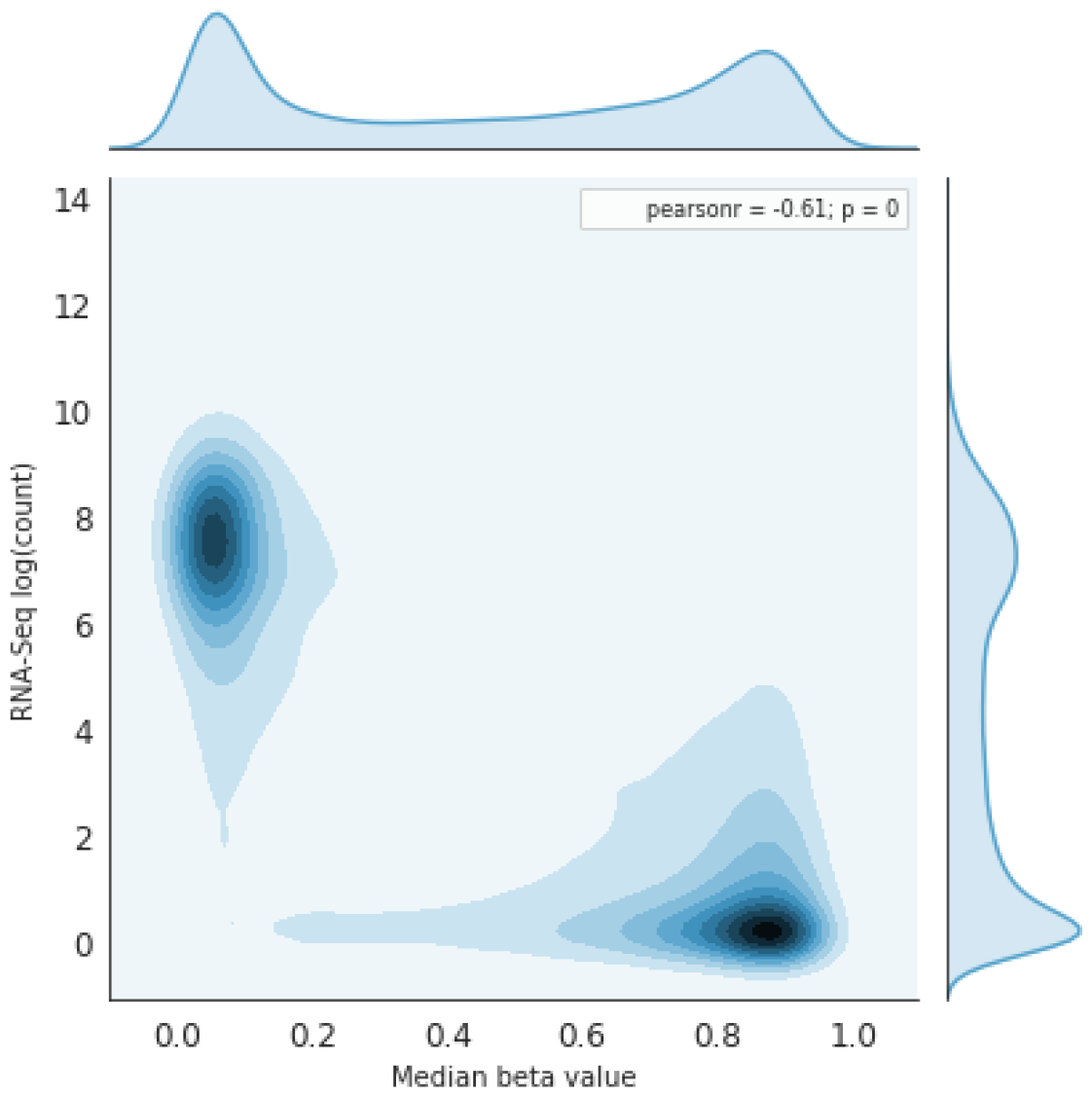
DNAm vs. RNA exp. Joint kernel density

**Figure 5.**
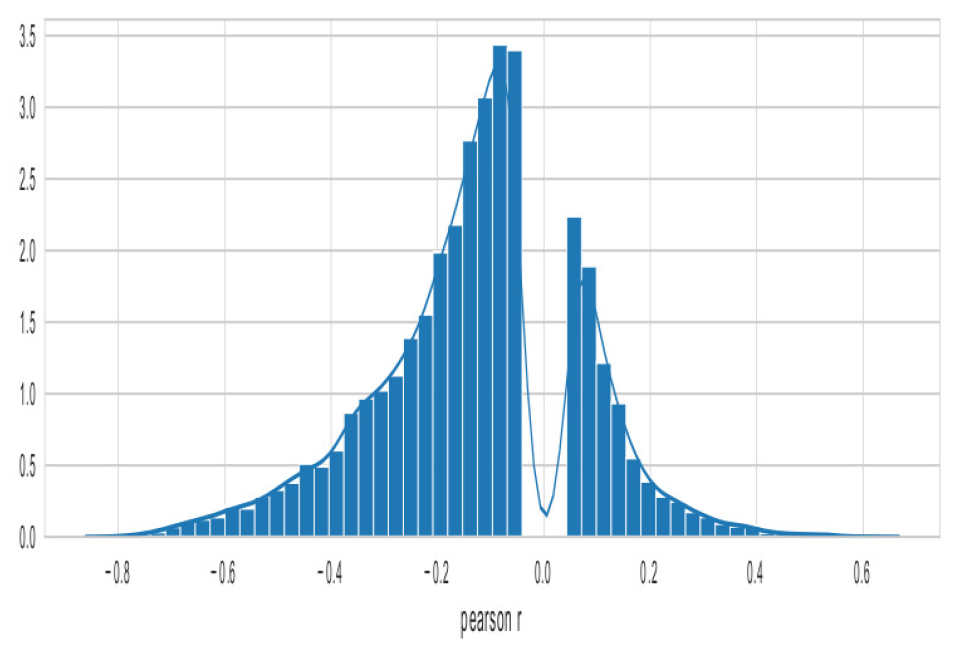
Pearson r correlations of DNAm & RNA expression

**Figure 6.**
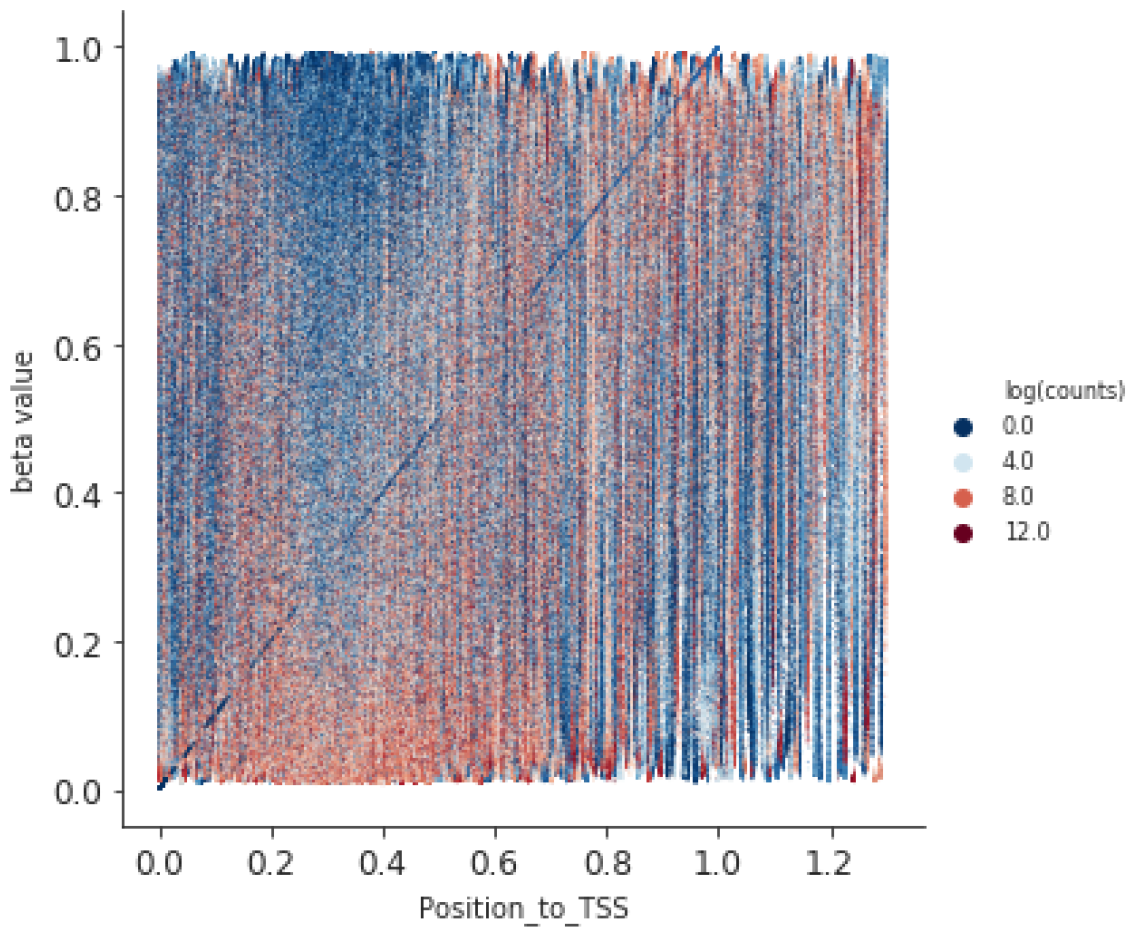
DNA Methylation & RNA expression

**Figure 7.**
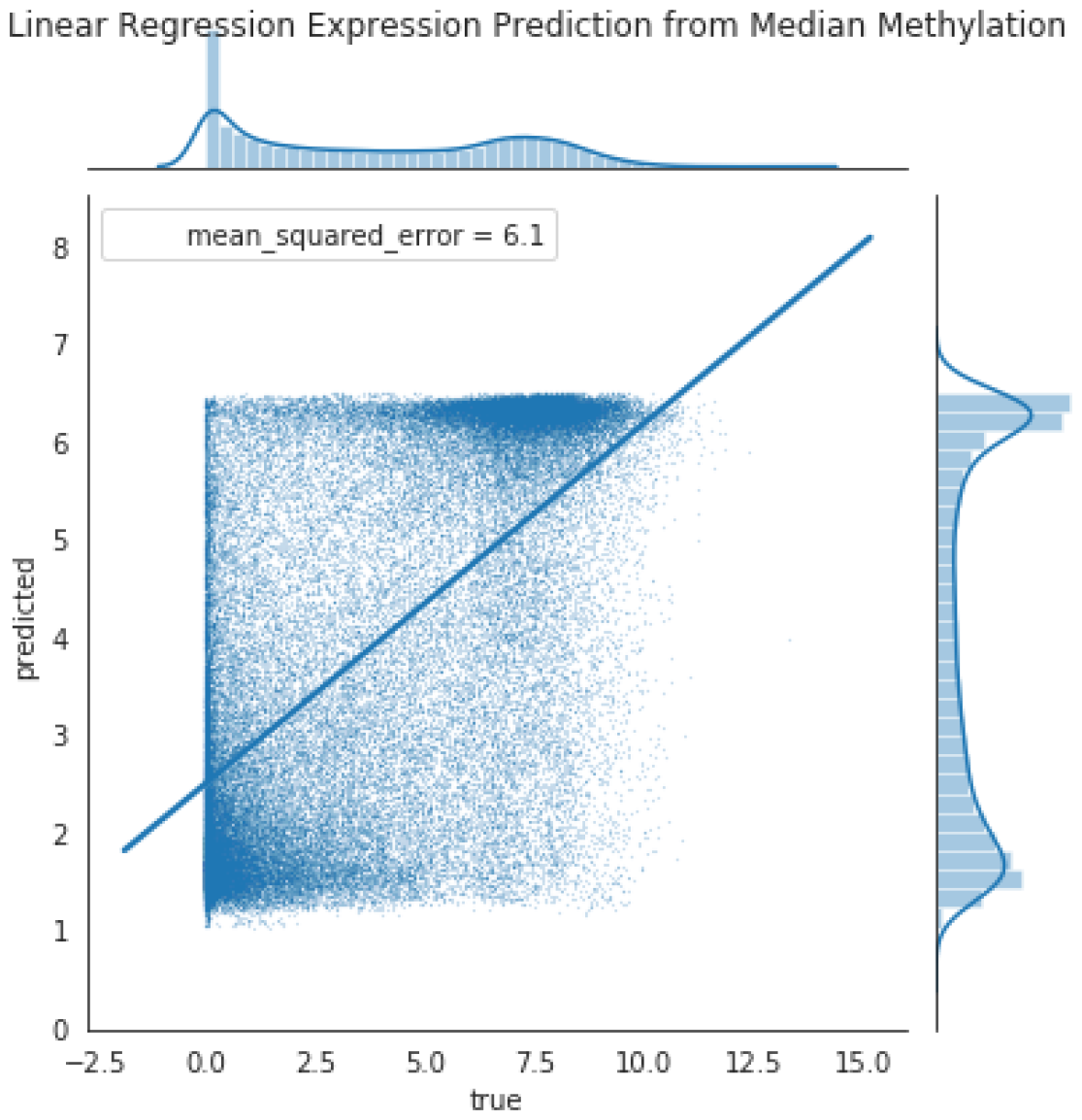
DNAm predicts RNA_exp_ via Lin. Reg.

**Figure 8.**
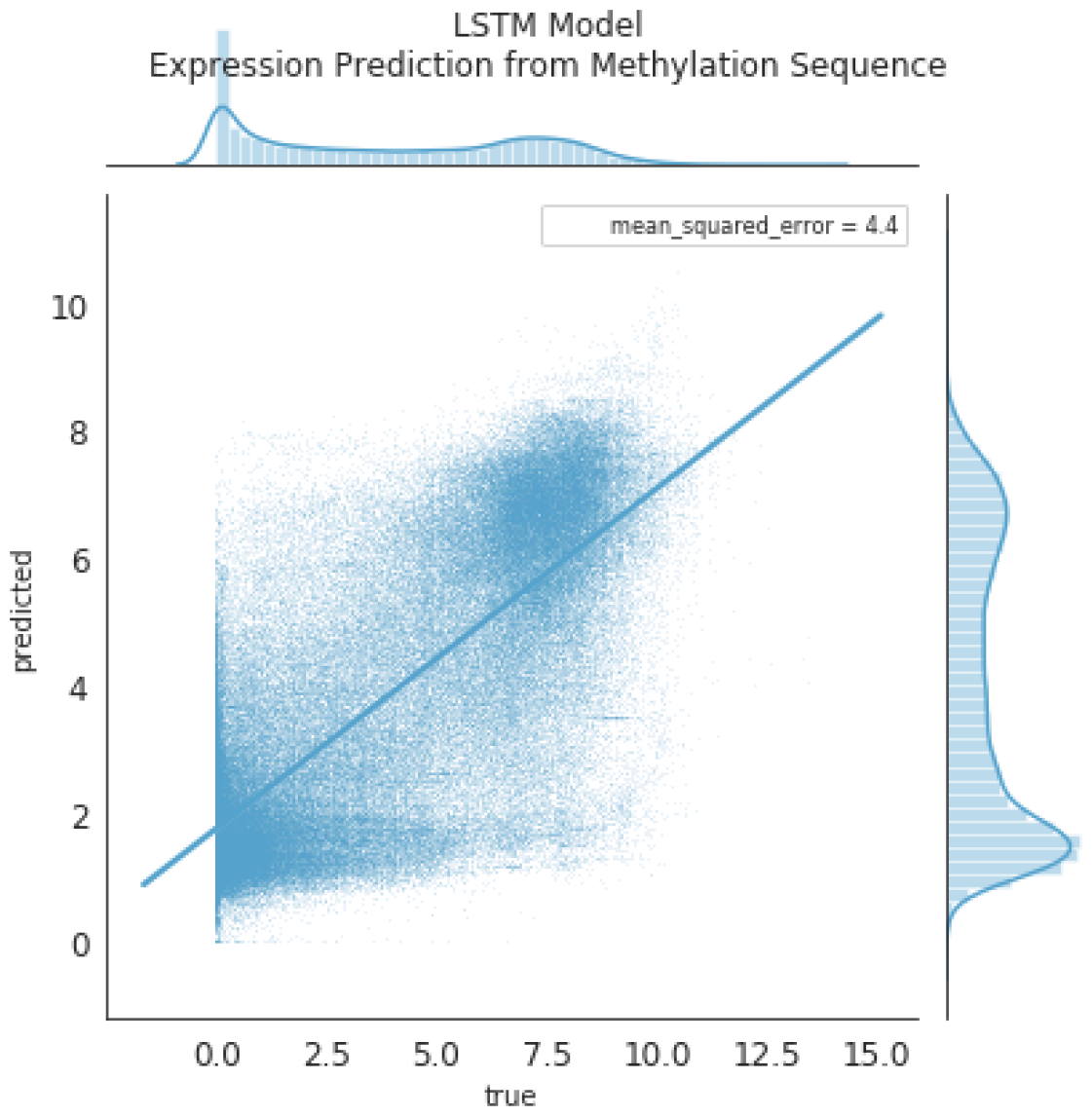
DNAm predicts RNA_exp_ via LSTM

## 5 Conclusion

The `autogdc` package improves developer workflows by integrating the querying, downloading, organization, and analysis functions of a bioinformatics study into one simple package. Furthermore, the inclusion of several machine learning tools for assessing the relationship between DNA methylation and transcription will facilitate research into epigenetic regulatory mechanisms.

## 5.1.2 Contributions

C.A.B is developer and writer, J.D.W. is PI

